# High expression of *VRT2* increases the number of rudimentary basal spikelets in wheat

**DOI:** 10.1101/2021.08.03.454952

**Authors:** Anna E. Backhaus, Ashleigh Lister, Melissa Tomkins, Nikolai M. Adamski, James Simmonds, Iain Macaulay, Richard J. Morris, Wilfried Haerty, Cristobal Uauy

## Abstract

Spikelets are the fundamental building blocks of *Poaceae* inflorescences and their development and branching patterns determine the various inflorescence architectures and grain yield of grasses. In wheat, the central spikelets produce the most and largest grains, while spikelet size gradually decreases acro- and basipetally, giving rise to the characteristic lanceolate shape of wheat spikes. The acropetal gradient correlates with the developmental age of spikelets, however the basal spikelets are developed first and the cause of their small size and rudimentary development is unclear. Here, we adapted G&T-seq, a low-input transcriptomics approach, to characterise gene expression profiles within spatial sections of individual spikes before and after the establishment of the lanceolate shape. We observed larger differences in gene expression profiles between the apical, central and basal sections of a single spike than between any section belonging to consecutive developmental timepoints. We found that *SVP* MADS-box transcription factors, including *VRT-A2*, are expressed highest in the basal section of the wheat spike and display the opposite expression gradient to flowering E-class *SEP1* genes. Based on multi-year field trials and transgenic lines, we show that higher expression of *VRT-A2* in the basal sections of the spike is associated with increased numbers of rudimentary basal spikelets. Our results, supported by computational modelling, suggest that the delayed transition of basal spikelets from vegetative to floral developmental programmes results in the lanceolate shape of wheat spikes. This study highlights the value of spatially resolved transcriptomics to gain new insights into developmental genetics pathways of grass inflorescences.

**One sentence summary:** Large transcriptional gradients exist *within* a wheat spike and are associated with rudimentary basal spikelet development, resulting in the characteristic lanceolate shape of wheat spikes.

## Introduction

The arrangement of flowers in individual plants of the same species is highly conserved and follows a systematic and rhythmic pattern. This systematic appearance of flowers is not surprising, as floral architectures are determined by the regular initiation of flower primordia on the flanks of the apical meristem and their rate of initiation and developmental fate are under strong genetic control (Prusinkiewicz et al., 2007). The unifying feature of floral architecture in grasses (*Poaceae*) is the formation of all flowers (termed florets) within spikelets (Kellogg et al., 2013). Spikelets are the fundamental building blocks of grass inflorescences and their development and branching patterns determine the various inflorescence architectures of grasses (e.g., spikes, panicles). Wheat (*Triticum aestivum*) forms a spike shaped inflorescence, in which sessile spikelets are directly attached to the inflorescence axis (or rachis) in a distichous phyllotaxis (Koppolu and Schnurbusch, 2019). Upon floral transition, the vegetative meristem ceases to initiate leaf primordia and transitions into the inflorescence meristem (IM). During the Double Ridge stage (DR) of wheat spike development, the IM initiates a lower leaf ridge and an upper spikelet ridge (or primordia) during each iteration. Within the inflorescence the upper ridges differentiate into spikelet meristems, while the lower ridges are suppressed upon flowering (Bommert and Whipple, 2018). DR initiation will continue at the IM until the terminal spikelet stage, when IM forms a final spikelet (Koppolu and Schnurbusch, 2019). Spikelet initiation and development has been extensively studied in wheat and other monocot crops, such as rice (*Oryza sativa*), maize (*Zea mays*), and barley (*Hordeum vulgare*), as the number of spikelets per spike is a major determining factor for grain number and thus yield per spike.

Not all spikelets across the wheat spike, however, produce the same amount of grain. The central spikelets produce the most and largest grains, while spikelet size gradually decreases acro- and basipetally. Within a single spike, the most apical and basal spikelets might produce no or only one grain while the central spikelets of the same spike set 3-5 grains. Bonnett (1966) documented that this distinct lanceolate shape of the wheat spike is first established during the Glume Primordia (GP) stage (just after the DR stage). This asynchronous development among the spikelets is maintained throughout the development of the spike. The gradual decrease in spikelet size from the central to apical section of the spike can be explained by the continuous development of new spikelet ridges from the apical inflorescence meristem: the most apical spikelets are the youngest and had the least time to develop. However, basal spikelets are initiated first and it is unclear why they remain smaller than their central counterparts. In the mature spike the most basal one or two spikelets are often only formed in a rudimentary manner, with small glumes present but all floral structures remaining immature.

Efforts to understand the genetics of wheat spikelet initiation and development have focused on members of the MADS-box transcription factor (TF) family, which play central roles in the flowering gene models (Zhao et al., 2006). Li et al. (2019) showed that MADS-box genes of the *SQUAMOSA-clade, VERNALISATION1 (VRN1), FRUITFULL2 (FUL2)* and *FRUITFULL3* (*FUL3*), have overlapping functions in controlling the timing of the transitions from the vegetative to IM as well as the formation of the terminal spikelet. In *vrn1ful2*-null mutants, the IM remained indeterminate causing the mutants to form more spikelets per spike. However, all lateral spikelets were replaced by leafy shoots in the *vrn1ful2* double and *vrn1ful2ful3* triple mutants (Li et al., 2019). These mutants had increased expression of genes belonging to the *SHORT VEGETATIVE PHASE* (*SVP*) family of MADS-box genes, including *VEGETATIVE TO REPRODUCTIVE TRANSITION 2* (*VRT2*). Subsequent studies determined that overexpression of *VRT2* led to reversion of basal spikelets to spikes and the downregulation of other MADS-box genes required for floral development, including members of the *SEPALLATA1* (*SEP1*) clade (Li et al., 2021). Together, these studies exemplify the importance of the temporal sequence of flowering gene expression for the correct development of the wheat spike.

Attempts to unravel the genetic network controlling wheat spike development have focused on these temporal changes in expression patterns across consecutive developmental stages. For example, Li et al. (2018) and Feng et al. (2017) performed transcriptome profiling using pooled samples of multiple complete spikes from six (vegetative to floret differentiation) and four (double ridge to young floret) developmental stages, respectively. In a few cases, studies have examined the expression patterns of individual genes (via quantitative reverse transcription (qRT)-PCR) and found gene expression gradients along the spike. For example, Debernardi et al. (2017) demonstrated that *APETALA2* (*AP2*) is expressed higher in the apical section of wheat spikes than in central or basal sections. This *AP2* expression gradient was associated with morphological changes along the same spike. This study alongside work in barley (Youssef et al., 2017), suggests that gene expression gradients *within* individual developmental stages could be important to further unravel the genetic control of spike development. However, despite its potential biological significance, spatial transcriptome profiles along the spike have yet to be investigated in wheat.

In this study, we aimed to characterise gene expression profiles along the spike during the establishment of the lanceolate shape of the wheat spike from DR to GP. We adapted G&T-seq (<u>G</u>enome and <u>T</u>ranscriptome sequencing), a low-input sequencing approach to sequence the transcriptome of the sections. Recently, Giolai et al. (2019) adapted the protocol to identify expression differences across single leaves of *Arabidopsis* (GaST-seq), demonstrating that the G&T-seq method can be readily used for sequencing of hand harvested, small input plant material without the need of previous tissue dissociation or treatment. G&T-seq is thus comparable to methods using laser-micro dissection followed by sequencing to achieve spatially resolved transcriptome wide sequencing data. In comparison, the available transcriptome sequencing methods at higher resolutions (such as single cell RNA-seq or fluorescence-activated cell sorting (FACS)) are not spatially resolved as the complete tissue is dissolved into single cells for barcoding or selection prior to sequencing (Rich-Griffin et al., 2020).

We sequenced the apical, central, and basal sections of individual spikes before (DR) and after (GP) the establishment of the lanceolate shape. Gene expression profiles differed most strongly between spatial sections of the same spike, as opposed to temporal sections (any two sections from different timepoints). Members of the *SVP* gene family were expressed most highly in the basal sections with expression decreasing upwards from the base (acropetally), while members of the *SEP1* gene family showed the opposite expression pattern, i.e. most highly expressed in apical sections, with expression decreasing towards the base (basipetally). The increased number of rudimentary basal spikelets due to *VRT-A2* misexpression supports the hypothesis that high expression levels of *SVPs* in the basal section delays spikelet establishment, leading to their rudimentary shape in the mature spike. This study highlights that spikelets within the same spike experience significantly different flowering signals due to their consecutive development and spatial position within the spike. Acknowledging these differences can help us gain a better understanding of the genetic flowering pathway of grass inflorescences.

## Results

### Low-input sequencing enables spatial analysis of the wheat spike transcriptome

To investigate transcriptional differences between the apical, central, and basal section of developing wheat spikes, we adapted the low-input G&T sequencing (G&T-seq) method for RNA-seq of small plant tissue sections. G&T has been developed for single-cell RNA and DNA sequencing of mammalian systems (Macaulay et al., 2015) and was previously adapted for *Arabidopsis thaliana* (GaST-seq; Giolai et al., 2019). We collected four individual developing wheat (cv Paragon) spikes at both the double ridge (DR) and glume primordia (GP) stage and hand-dissected them into apical, central, and basal sections (Figure 1A).

**Figure 1:**
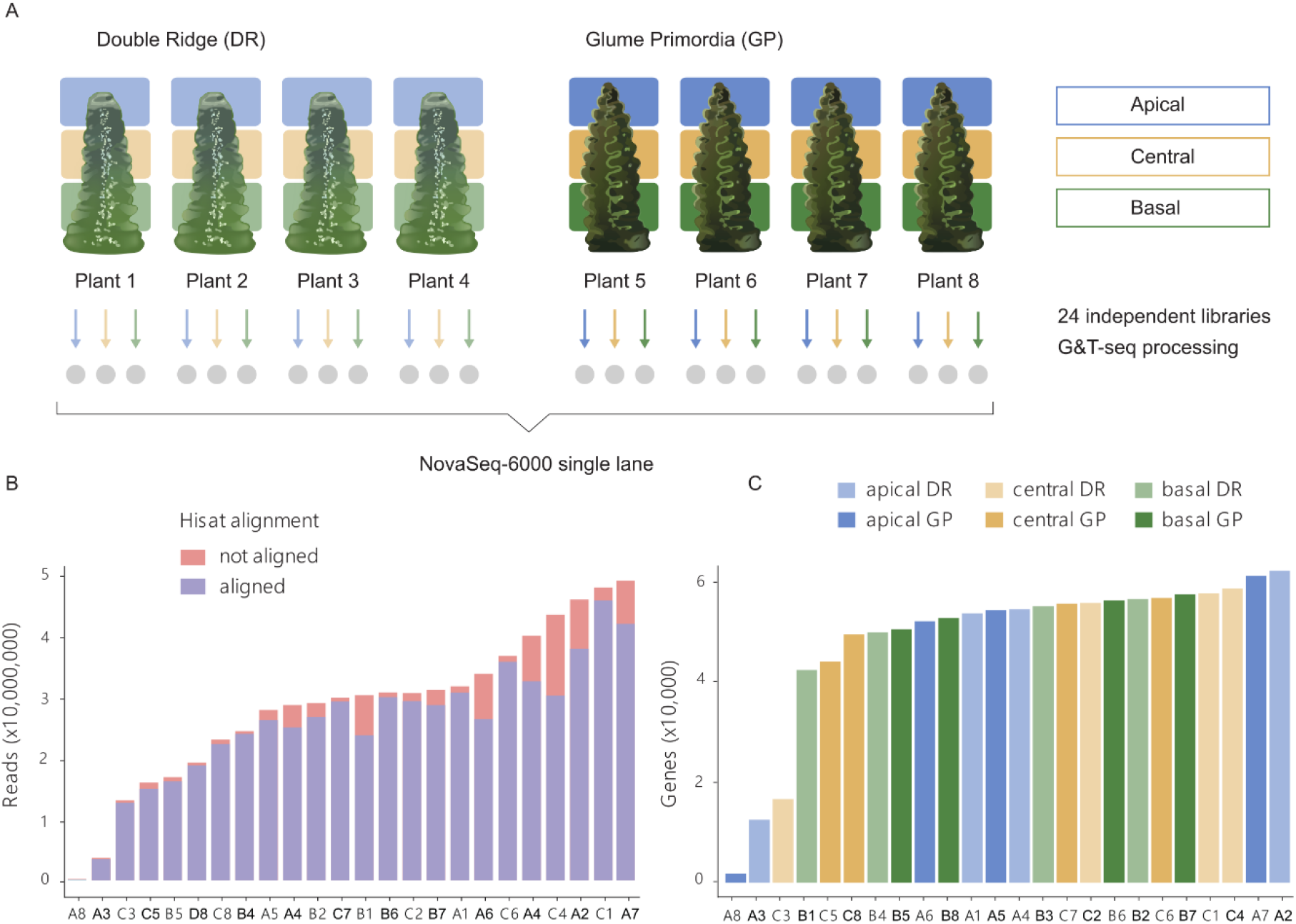
(A) Summary diagram of low-input G&T tissue collection and sequencing approach. Grey circles indicate the 24 individual libraries prepared for sequencing from each individual tissue section dissected from individual spikes. (B) Reads per library after trimming and quality controls (see Methods). Stacked bars indicate the number of reads aligned (blue) and not aligned (red) by HISAT to the RefSeqv1.0 genome. (C) Number of expressed genes (>10 read counts) per library based on tissue section and Waddington developmental stage (DR: Double Ridge; GP: Glume Primordia). In (B and C), the X-axis indicates the ID of each sample which is composed of the tissue section (A: apical, C: central, B: basal) and plant number (1-8) as indicated in (A). Detailed quality control data for each library is provided in Supplemental Table S1.

On average, samples had 28,799,626 reads (coefficient of variation (CV) 43%), of which 90% (CV 8.5%) aligned to the genome post adaptor trimming (Figure 1B, Table 1). Furthermore, the number of aligned reads and the number of expressed genes per library was largely homogenous among the spatial sections and Waddington stages (Table 1). On average, 47,313 genes per library were expressed (>10 read counts) and we found no difference (*P* > 0.56) in the number of expressed genes across spatial (apical, central, basal) or between temporal (DR, GP) conditions (Figure 1C). We excluded three libraries with low average number of expressed genes (difference greater than five times the standard deviation; Figure 1C, Supplemental Table S1) and two libraries because they were strong outliers in the principal component analysis (PCA; Supplemental Figure S1A). In total, 19 RNA-seq libraries (DR: 3 apical, 4 central, 3 basal; GP: 2 apical, 3 central, 4 basal) passed our selection criteria and were used in the subsequent analyses. We identified 91,646 genes being expressed across these 19 libraries.

**Table 1:** Average number of reads aligned to the RefSeqv1.0 genome and expressed genes (>10 read counts) in the three tissue sections and two Waddington developmental stages (DR: Double Ridge; GP: Glume Primordia) (n = 4 biological replicates per tissue section * developmental stage).

|  | Reads aligned |  | Genes expressed |  |
| --- | --- | --- | --- | --- |
|  | DR | GP | DR | GP |
| **Apical** | 26,342,934 | 23,913,657 | 44,488 | 41,152 |
| **Central** | 29,823,023 | 25,854,540 | 45,913 | 50,074 |
| **Basal** | 25,174,482 | 23,627,494 | 49,513 | 52,740 |

### Transcriptome-wide differences are largest between the apical and basal sections of the spike

To investigate global differences among the 19 RNA-seq libraries, we performed a principal component analysis (PCA; Figure 2A). The first two PCs explained 19% and 16% of the overall variance present in the libraries. We observed that the two PCs separated libraries by the spatial position (apical, central, basal) rather than developmental stage (DR, GP). There was a clear separation between libraries originating from apical and basal spike sections, while libraries from central sections were dispersed between these two clusters (Figure 2A). We investigated PC1 to PC6 and found that none of these combinations clustered libraries by developmental stage (Supplemental Figure S1B). Given that we sequenced developing spike sections of single plants, as opposed to the more commonly employed pooling of multiple biological samples, we found as expected some degree of heterogeneity between samples from the same location and stage (Figure 2A).

**Figure 2:**
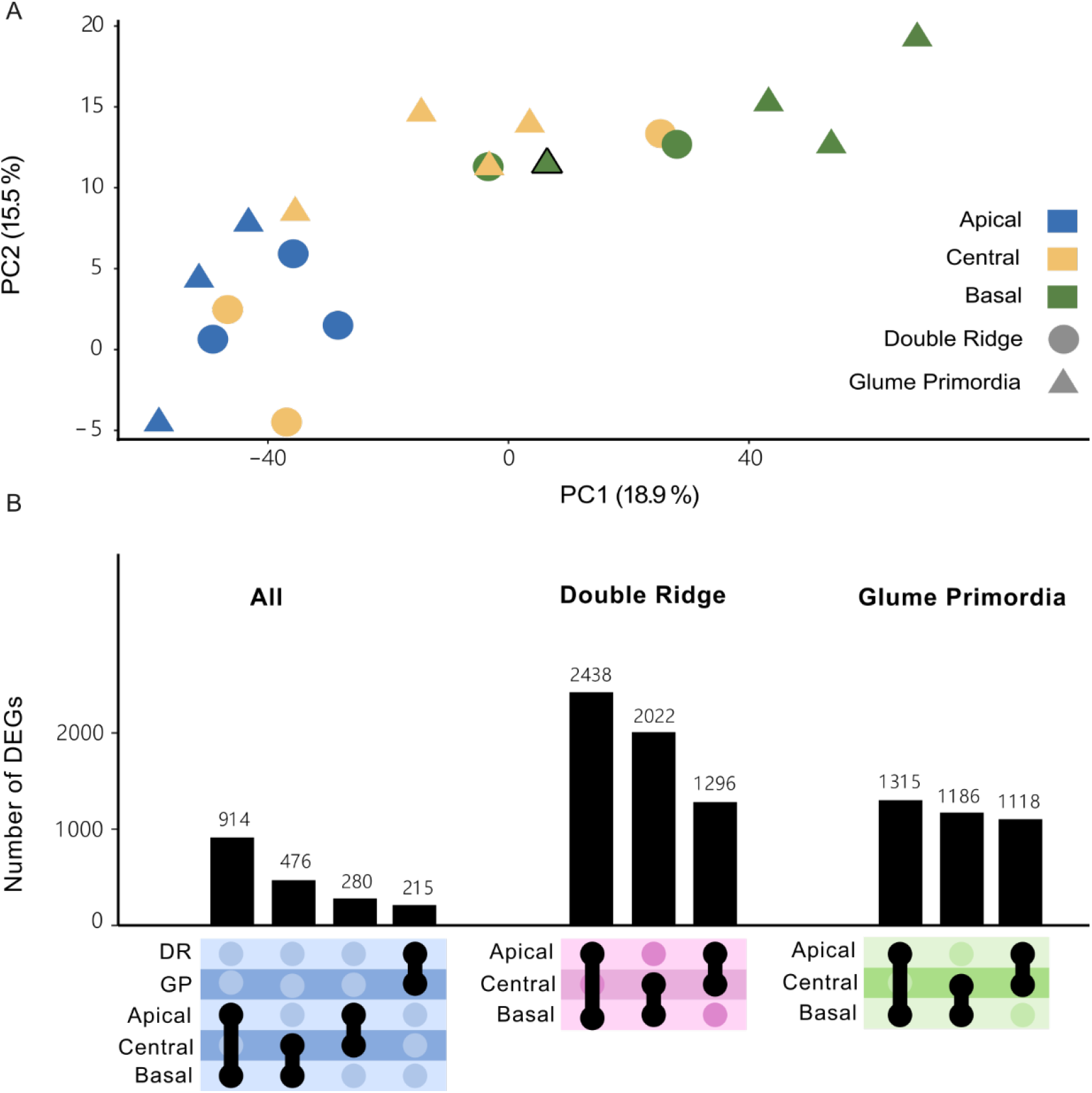
(A) Principal component analysis (PCA) on the 19 transcriptome libraries from apical (blue), central (yellow) and basal (green) sections of Double Ridge (circles) and Glume Primordia (triangles) spikes. Black bordered triangle is plant 8 (GP) in which the basal section clustered closer with central-GP sections than the other basal-GP sections. (B) UpSet plot showing the number of differentially expressed genes (DEGs) between spatial sections and Waddington stages. Black border indicates Plant 8, which is an outlier of basal GP replicates.

To investigate this variation further, we quantified changes in gene expression across biological replicates by calculating CVs for each gene (see Methods). The median CV for a gene across the biological replicates was 39% (Supplemental Figure S2) with a Q1-Q3 interquartile range between 24% and 62%. We also calculated the CV per gene for published datasets from Li et al (2018) and Feng et al (2017). Both studies sequenced developing wheat spikes at similar developmental stages, pooling many spikes per sample. Li et al. (2018) pooled between 100 to 200 spikes of winter wheat (KN9204) per sample, while Feng et al. (2017) reported pooling of 10 to 50 spikes (cv. Chinese Spring) per sample. In both studies the median CV of a gene was lower (14% and 21%, respectively) than in our study. The larger CVs in our data could be explained by the biological variation that exists between individual plants, which may have been reduced by the pooling of many spikes in both Li et al. (2018) and Feng et al. (2017).

We first analysed differentially expressed genes (DEGs) between the DR and GP stage and between apical, central and basal sections across the two Waddington developmental stages. The number of DEGs between DR and GP (215 genes) was smaller than the number of DEGs identified between the spatial positions, which ranged from 280 DEGs between central and apical sections to 914 DEGs between the apical and basal sections (Figure 2B). Next, we compared the apical, basal and central sections within each Waddington developmental stage. We identified more DEGs by comparing the spatial sections within either Waddington stage individually than in the combined analysis. The number of DEGs between apical and basal sections at each stage (DR: 2,438; GP: 1,315) were similar to the number of DEGs between central and basal sections (DR: 2,022; GP: 1,186). The number of DEGs between these sections at DR, however, was nearly double the number of DEGs at GP. In contrast, the number of DEGs between apical and central sections was similar at both stages (DR: 1,296; GP: 1,118), suggesting that the basal section of the spike is most different in the earlier developmental stage. Only 11% of the DEGs were shared between DR and GP in the apical to basal comparison, 7% between the central to basal DEGs, and 5% between apical to central DEGs. In total, we identified 5,353 unique genes as differentially expressed between any of the three sections at either Waddington stage (Supplemental Table S2). Overall, the number of DEGs was largest between the apical and basal sections, reflecting the strong spatial clustering observed in the PCA graph, but most genes that were differentially expressed across the spike did not maintain this gradient over the two developmental stages. In summary, despite the high biological variation in gene expression in our data compared to previous pooled whole-spike studies, we could detect transcriptome wide differences between the spatial sections of developing wheat spikes.

### The *SVP* MADS-box transcription factors have opposing expression profiles to flowering E-class genes

To further investigate the differences in expression across the spike and to identify genes with similar expression patterns, we performed hierarchical and k-means clustering. We restricted the clustering to the 5,353 genes identified as differentially expressed across the spike at either one or both Waddington stages (Figure 3A). We identified seven non-redundant clusters, each containing between 8% to 21% of the 5,353 DEGs (Figure 3A, Supplemental Figure 3A). Both hierarchical and k-means clustering produced highly similar results (Supplemental Figure S3B). We identified 1,894 genes (35% of DEGs) to be more highly expressed in the apical section, either across both timepoints (503 genes, cluster 1), or only at DR (751 genes, cluster 7) or GP (640 genes, cluster 3). In the central section, 1,362 genes (25%) had higher relative expression at either DR (917 DEGs, Cluster 5) or GP (445 DEGs, Cluster 6). In the basal section, we observed 2,097 genes (39%) being more highly expressed. Cluster 4 contained the most DEGs (1,170) and was characterized by an upregulation of expression in the basal section at both Waddington stages, although this upregulation was higher at the DR stage. Another 927 genes were upregulated in the basal section, but only at the GP stage (cluster 2).

**Figure 3:**
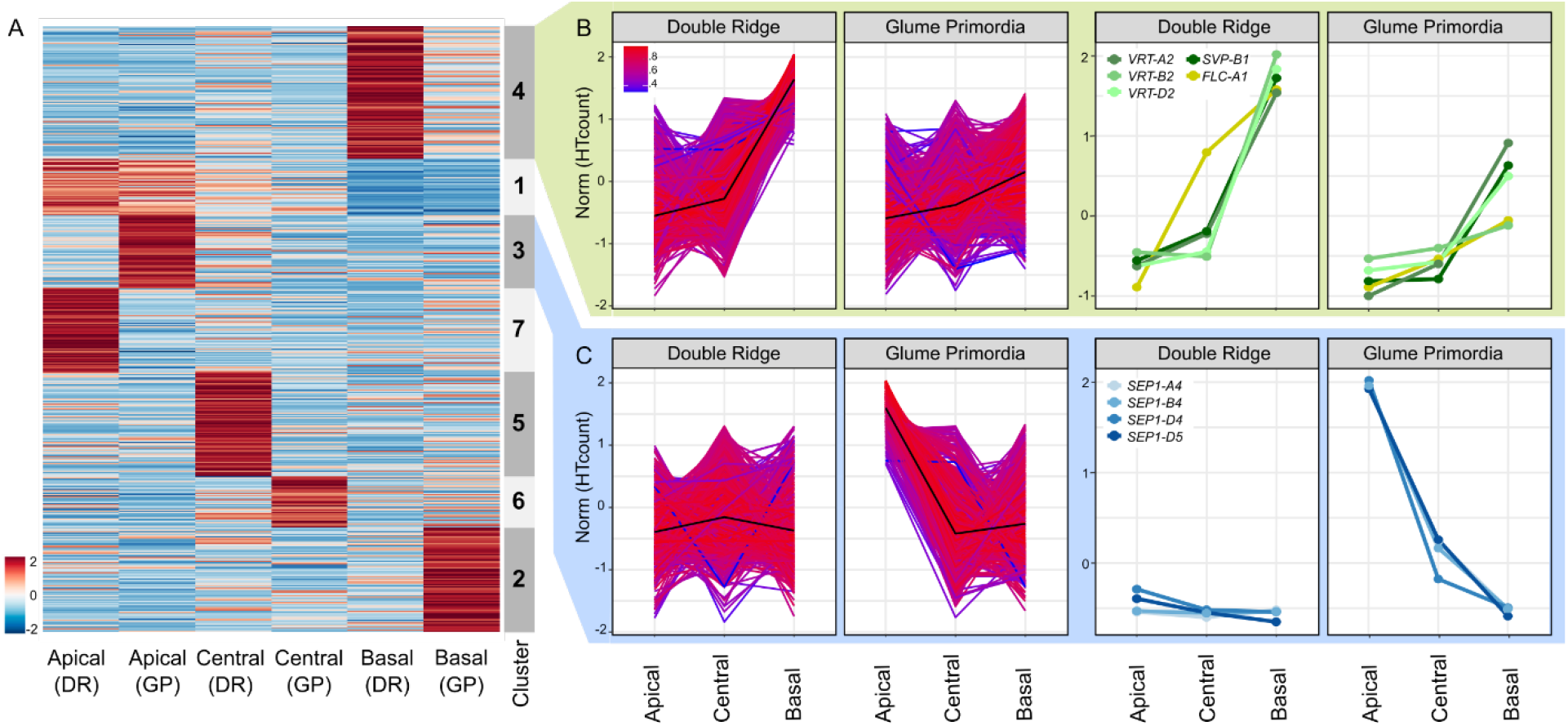
(A) Normalized expression matrix and K-means clustering of the 5,353 genes differentially expressed across the spike at either one or both Waddington stages. Colours (blue to red) show relative log2 expression of genes after normalisation. (B) Expression pattern of the 1,170 genes allocated to Cluster 4 (left), and of MADS-box transcription factors in the same cluster (right). Colours indicate how well the gene expression pattern fits the average expression pattern (black line). Red = best fit, Blue = least good fit. (C) Expression pattern of the 640 genes and MADS-box transcription factors of Cluster 3 as arranged in B. Norm = normalized and scaled gene expression. RefSeq1.1 gene IDs and raw expression values of genes shown in the right-hand panels are presented in Supplemental Table S2.

To further characterize the clusters, we independently tested for enrichment of TF families and gene ontology (GO)-terms (all GO-terms and TF families in Supplemental Table S3 and S4, respectively). Genes that were more highly expressed in the apical section were enriched for the GO terms “reproductive structure development” (GO:0048608) and “floral organ development” (GO:0048437; cluster 1) as well as for HD-Zip_IV and SRS TF families (*P* < 0.0001; cluster 1). In cluster 3 (highly expressed in the apical section at GP; Figure 3B) we found no significant enrichment of GO-terms (*P* < 0.03), but a significant enrichment of MADS_II TFs (*P* = 0.013). For clusters defined by an increased expression in the central sections (clusters 5 and 6) we detected an enrichment for GO-terms related to polyphosphate processes (GO:0006797/0006779), but a significant enrichment of the C2C2_CO-like TFs at DR (*P* = 0.036; cluster 5). Genes with higher expression in the basal section of the spike (cluster 4; Figure 3C) were enriched for a number of GO terms related to photosynthesis, (e.g. GO:0015979) and “negative regulation of flower development” (GO:0009910), as well as for MADS_II TFs (*P* = 0.084). Cluster 2, which was characterized by higher expression in the basal section only at GP, was enriched for “Jasmonic acid response” (GO:0009753) and Tify and C2C2_CO-like TFs (*P* < 0.04).

We were interested in further characterizing the expression patterns of the MADS-box TFs as they were significantly enriched in two of the seven clusters and are important in floral transition and development (Becker and Theißen, 2003; Feng et al., 2017). We detected 14 differentially expressed MADS-box TFs in our study, five of which were more highly expressed in the apical section, four of these only at GP stage (cluster 3) and one being consistently expressed across both Waddington stages (cluster 1). In contrast, five MADS-box TFs were more highly expressed in the basal section at both Waddington stages (cluster 4) and another two were more highly expressed in the basal section only at GP (cluster 2). An additional two MADS-box genes were part of the remaining clusters (Supplemental Table S4).

In the apical/GP cluster 3 we noticed that all MADS-box genes belonged to the *Triticum aestivum SEPALLATA1 (TaSEP1)* group (Figure 3C). All three homoeologs of *TaSEP1-4* (*TraesCS7A02G122000, TraesCS7B02G020800, TraesCS7D02G120500*) and the D-genome copy of *TaSEP1-5* (*TraesCS7D02G120600*) were part of this cluster. The *SEP* genes were expressed at relatively low levels at DR (Supplemental Table S2), but were significantly upregulated at GP, with their transcript levels being highest in the apical section. The increased expression of *TaSEP1-4* at GP was in agreement with their previously reported expression patterns in tetraploid wheat by Li et al. (2021).

In the contrasting cluster 4 (upregulation in basal sections), we noticed the presence of multiple MADS-box genes belonging to the *SVP* family (Figure 3B, right-hand panel), which consists of three genes in wheat (*SVP1, VRT2* and *SVP3*). Members of this family are important for the transition from vegetative to floral meristem identity in cereals (Trevaskis et al., 2007). All three homoeologs of *VRT2 (TraesCS7A02G175200, TraesCS7B02G080300, TraesCS7D02G176700*) and the B-genome copy of *SVP1* (*TraesCS6B02G343900*) were present in cluster 4. The cluster also contained *TaFLC-A1* (*TraesCS7A02G260900*), although it was expressed higher in DR-central sections compared to the *SVPs* and had a linear expression gradient at GP. All *SVPs* had very similar expression patterns, being strongly expressed in basal sections only. Expression of *SVPs* was higher in all DR sections compared to the equivalent section in GP. Constitutive over-expression of *SVP*-family members in wheat and barley has been shown to delay or even reverse floral development (Trevaskis et al., 2007; Li et al., 2021). This led to hypothesis that the rudimentary development of basal spikelets was associated with an increase in *VRT2* expression levels.

### *SVP* expression is higher in basal and peduncle sections and increased across all sections in *T. polonicum VRT-A2b isogenic lines*

To validate the expression pattern of *VRT2* in the individual spike analysis, we performed qRT-PCR on independently collected, pooled spike sections from cv Paragon, carrying the wildtype *VRT-A2a* allele (Figure 4A, blue curves). We included a later timepoint, Terminal Spikelet (TS), which is about 10 days after GP to study how *VRT2* expression changes in later stages. At TS stage, the central spikelets have developed multiple florets primordia. We also included a small part of the peduncle (stem) section just below the spike as an additional spatial section. We focused the expression analysis on the A-genome homoeolog, *VRT-A2*, as its role in spike, glume and grain development of wheat was recently characterised (Adamski et al., 2021; Liu et al., 2021).

**Figure 4:**
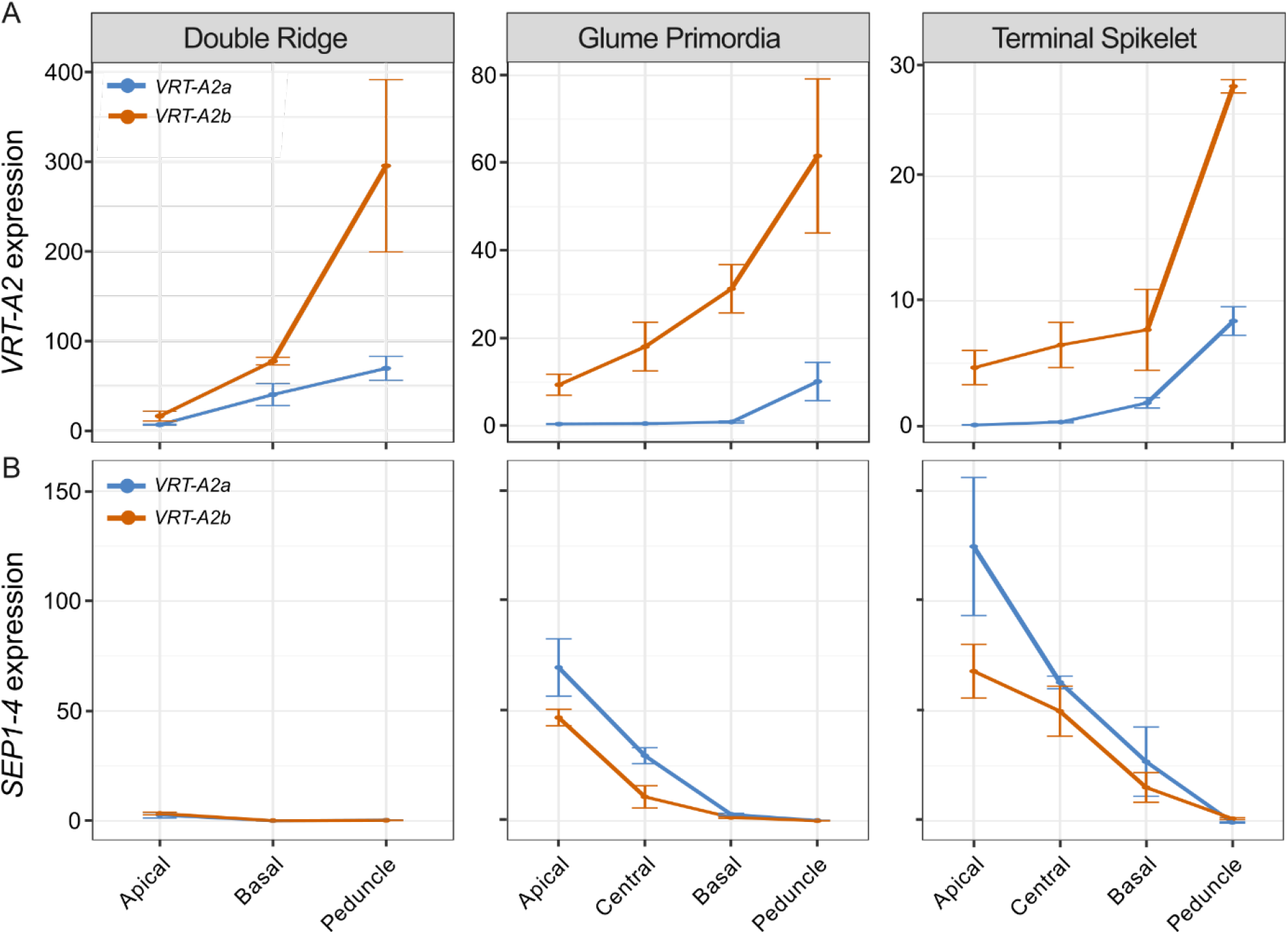
Relative expression (2^ddCT^) of *VRT-A2* (A) and *SEP1-4* (B) in the different sections of the spike across three timepoints in near isogenic lines (NILs) carrying either the wildtype *VRT-A2a* (blue) or the *VRT-A2b* allele from *Triticum turgidum* ssp. *polonicum* (orange). The data are shown as mean ± SE of gene expression compared with control gene Actin. N = 3 biological replicates (See Supplemental Table S5 for expression data and Supplemental Table S6 for statistical analysis of gene expression differences).

We identified a significant interaction effect between Waddington stage and spatial section; we thus analysed the three Waddington stages separately (Figure 4). At DR, we were limited to dissecting the spike into apical, basal and peduncle sections, as the small size of the spike meristem did not allow precise dissection of the central section when using multiple (pooled) spikes. At DR, we found *VRT-A2* marginally expressed, with significantly lower expression levels in the apical section compared to the basal (*P* = 0.003) and peduncle sections (*P* = 0.001). Although expression in the peduncle was higher than in the basal section at DR, this was not significant (*P* = 0.116). At GP, *VRT-A2* expression was borderline detectable in the apical, central and basal sections, but expression was significantly higher in the peduncle with respect to the three spike tissues (*P* < 0.001 for all three comparisons). Lastly, *VRT-A2* expression at TS stage was just detectable and significantly different between all sections (*P* = 6.6E-06). Overall, expression decreased significantly from DR to GP/TS Waddington stages in the apical (*P* = 0.00015), basal (*P* = 0.0074), and peduncle (*P* = 0.012) sections consistent with the previously reported strong downregulation of *VRT-A2* in the early wheat spike development (Li et al., 2021; Adamski et al., 2021; Liu et al., 2021). This is also consistent with the observed downregulation of *VRT-A2* orthologs upon floral transition in barley (Trevaskis et al., 2007) and rice (Harrop et al., 2018). As observed in the low-input RNA-seq data, the qRT-PCR data confirmed the strong basipetal gradient in *VRT-A2* expression across the spike at DR and revealed that its expression was even higher within the peduncle.

We hypothesised that the higher expression in the basal section of the wheat spike compared to the central and apical sections is associated with the rudimentary development of the basal spikelets. To test the effect of higher *VRT-A2* expression on basal spikelet development, we analysed the effect of the *Triticum turgidum* ssp. *polonicum VRT-A2b* allele on the expression gradient of *VRT-A2* and spike morphology. Adamski et al. (2021) showed that *VRT-A2* in *T. polonicum*, a tetraploid subspecies of wheat, carries a sequence re-arrangement in its first intron. This results in the higher expression of the *T. polonicum VRT-A2b* allele, with respect to the wildtype *VRT-A2a* allele, during early spike development. We performed qRT-PCR on a cv Paragon NIL carrying the *VRT-A2b* allele and compared *VRT-A2* expression against the Paragon wildtype NIL described above (Figure 4). Consistent with the results of Adamski et al. (2021), we detected significantly higher expression of *VRT-A2b* compared to the wildtype allele across most of the tissue sections (see Supplemental Table S6 for individual comparisons), and a progressive decrease in *VRT-A2b* expression over time (*P* = 0.031). In contrast to the wildtype NILs, ANOVA did not identify a significant interaction effect between spatial section and Waddington stage in *VRT-A2b* NILs (*P* = 0.18). We thus examined the overall expression patterns and found that across all three developmental stages *VRT-A2b* expression differences were significant (*P* < 0.0001). These results suggest that the basipetal expression gradient in the spike is maintained in the NILs with the *T. polonicum VRT-A2b* allele.

We also tested the effect of *VRT-A2* expression levels on *SEP1* expression in the tissue sections of *VRT-A2* NILs. We confirmed that *SEP1-4* expression is only marginally detectable at DR and differences in expression between the apical, basal, and peduncle section are hardly detectable with only marginally higher expression in the apical sections (*P* = 0.015) at this stage. At GP, *SEP1-4* expression is significantly higher towards the tip of the spike (*P* < 0.0001) consistent with the low-input RNA-seq data. Furthermore, *SEP1-4* expression was significantly lower in *VRT-A2b* NILs compared to the wildtype allele across all spike sections (*P* = 0.008), confirming that higher *VRT-A2* expression can negatively affect *SEP1-4* expression. Similar trends were observed at TS, where expression was significantly lower in basal sections (*P* < 0.0001) and the gradient across the spike was maintained in *VRT-A2b* NILs, but expression was overall lower (*P* < 0.003). *SEP1-4* is not expected to be expressed in vegetative tissue such as the peduncle, therefore the lack of expression in this tissue across the three stages indicates that no floral tissue was accidently sampled as peduncle (Fig. 4B).

### Misexpression of *VRT-A2b* in *T. polonicum* increases rudimentary basal spikelet numbers

To evaluate if the higher expression of *VRT-A2* in basal spikelets affects their development, we examined the *VRT-A2* NILs (BC_4_ and BC_6_) sown as winter crops in four environments. In each field trial, we evaluated the number of rudimentary basal spikelets (RBS), that is spikelets which are reduced in size and do not contain mature grains (Figure 5A). The number of RBS was significantly increased in NILs carrying the *VRT-A2b* allele in all four environments (*P* < 0.0001, except Morley 2017 *P* < 0.01; Figure 5B; Supplemental Table S7). The *VRT-A2a* NILs and the recurrent parent Paragon had on average 1.85 RBS, whereas *VRT-A2b* NILs produced on average 2.91 RBS. A similar difference in RBS between the NILs was observed in glasshouse conditions (*VRT-A2b* effect of+1.6 RBS; Supplemental Table S8; Figure 5C).

**Figure 5:**
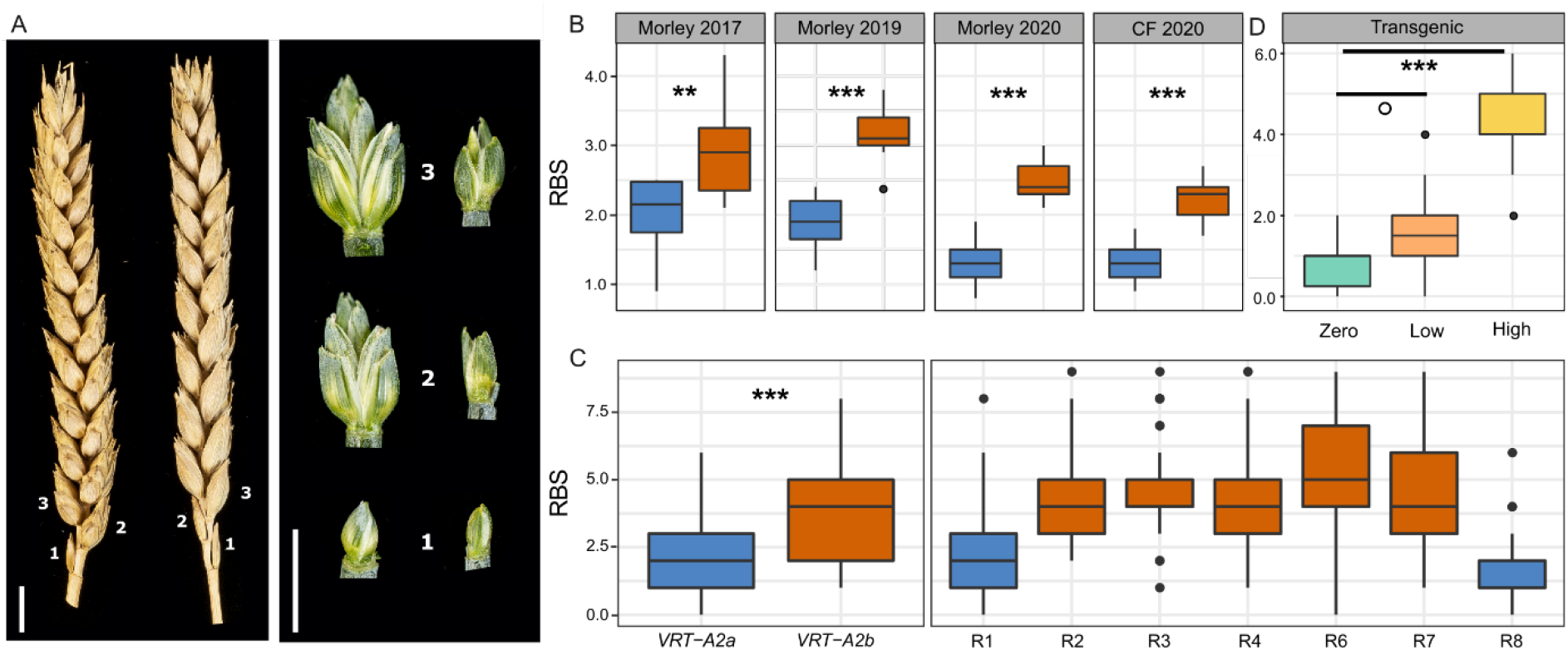
Phenotypic difference between *VRT-A2a* (blue) and *VRT-A2b* (orange) on rudimentary basal spikelet numbers (RBS). (A) Mature spikes from the field (left) and dissected basal spikelets at anthesis (right) from the glasshouse. Numbers indicate position along the spike starting at the base. Scale bar = 1 cm. (B) Number of RBS per spike from 10-ear samples collected in the field at maturity at Morley (2017, 2019 and 2020) and Church Farm (CF, 2020). (C) Number of RBS recorded in the glasshouse for the NILs (left panel) and for seven critical recombinant lines (R1-R4, R6-R8; see Supplemental Table S8 for graphical genotype of these lines from Adamski et al. (2021). (D) RBS per spike recorded for the transgenic lines carrying zero, low (1-5) or high (9-35) copy-number insertions of *VRT-A2b* in cv. Fielder. In B-D, the box represents the middle 50% of data with the borders of the box representing the 25th and 75th percentile. The horizontal line in the middle of the box represents the median. Whiskers represent the minimum and maximum values, unless a point exceeds 1.5 times the interquartile range in which case the whisker represents this value and values beyond this are plotted as single points (outliers). *P* values: o ≤0.1; ** ≤ 0.01; *** ≤ 0.001.

Furthermore, we also recorded the number of RBS in seven homozygous BC_6_ recombinant lines used to fine-map *VRT-A2b* by Adamski et al. (2021). The RBS phenotype was mapped in complete linkage with the 50.3 kbp interval containing *VRT-A2* (Figure 5C; Supplemental Table S8). This genetic and phenotypic data suggests that the increase in RBS is a pleiotropic effect of the *T. polonicum VRT-A2b* allele and supports the hypothesis that misexpression of *VRT-A2* negatively affects spikelet development in the base of the spike. In Paragon, the first (sometimes second) rudimentary basal spikelet fully develops the floral organs of florets one and two (e.g. lemma, palea, stamen, and ovary), however these are severely reduced in size and delayed in development compared to the florets of central spikelets just before flowering (~Waddington stage 8-10; Supplemental Figure S4). At this stage, the further growth and development of these basal florets is stopped and in the mature spike only the glumes of RBS are visible. In NILs carrying the *VRT-A2b* allele the development of the most basal spikelet is very similar to the wildtype. However, the second, third and sometimes fourth spikelet also display similar signs of reduced development, leading to the larger number of rudimentary basal spikelets (Figure 5B, C; Supplemental Figure S4).

To validate the phenotypic effect of *VRT-A2*, we analysed transgenic wheat lines transformed with the complete genomic *T. polonicum VRT-A2b* sequence (including the native promoter and the intron 1 re-arrangement). Transgenic T_1_ lines were classified based on the transgene copy number which was previously shown by Adamski et al. (2021) to be highly correlated with *VRT-A2* expression levels in multiple tissues. We phenotyped lines with zero (n = 2 independent events; 5 plants each), low (1-5 transgene copies; n = 4 independent events; 5 plants each) and high (9-35 transgene copies, n = 2 independent events; 5 plants each) transgene copy number. We identified a significant and stepwise increase in the number of RBS with transgenic copy number, from 0.8 RBS (zero copy) to 1.6 RBS (low copy; *P* = 0.078 vs zero copy) and to 4.3 RBS (high copy; *P* < 0.0001 vs zero copy) (Figure 5D; Supplemental Table S9). The low copy number lines had an average increase of 0.8 RBS with respect to the zero copy number lines, equivalent to the average difference between the *VRT-A2a* and *VRT-A2b* NILs in the field (*VRT2-A2b* effect of +1.1 RBS). The high copy number lines produced on average 4.3 RBS, which is higher than the *VRT-A2b* NILs and similar to the number of RBS observed in *T. polonicum* (3.75 ± 0.62 RBS; n = 16 spikes). The dosage-dependent effects observed in the transgenic lines provide further evidence that elevated expression of *VRT-A2* leads to increased number of rudimentary basal spikelets in polyploid wheat.

## DISCUSSION

### High-resolution spatial transcriptomics in crops

We hypothesised that the establishment of the lanceolate shape in wheat spikes could be manifested in gene expression differences between the apical, central and basal sections of a developing spike, as has been shown using qRT-PCR for individual genes in wheat (*AP2*; Debernardi (2017)) and barley (*VRS2*, Youssef et al. (2017)). However, currently available transcriptome data (e.g., Li et al. (2018) and Feng et al. (2017)) lack the spatial resolution *within* each individual developmental stage to answer this question. This focus on ‘between stage’ comparisons (as opposed to within a single stage) is perhaps related to the technical challenges of dissecting and sectioning young meristems. Given the relatively small size of these spike meristems (0.2 mm length at Transition Stage; 3 mm length at Terminal Spikelet stage), RNA-seq methods require bulking of multiple individuals (usually between 30 and 50 different plants) to accumulate enough tissue for a single RNA-seq sample. If one sought to further section each meristem, this would require even further bulking. While laborious, this is achievable; however, under this scenario, the challenge is to properly stage ~100 plants to an equivalent developmental stage. Furthermore, it can be technically challenging to section these young spikes each time into the exact same apical, central and basal sections. Consequently, the spatial resolution in gene expression within a wheat spike at individual developmental stages has remained largely uncharacterised to date.

To address this challenge, we adapted the G&T method for micro-scale spatial-transcriptomics workflow (Macaulay et al., 2015; Giolai et al., 2019), to conduct RNA-seq of the apical, central and basal sections of individual, hand-dissected wheat spikes. This highly-automated workflow requires low tissue input and allowed us to combine 24 Nextera libraries into a single Illumina NovaSeq lane. For 19 out of the 24 samples the method worked successfully, determined by >20,000 expressed genes per library and the clustering among biological replicates. We found that the number of expressed genes per library was on average similar to the number of genes reported for bulked whole spike RNA-seq samples (Feng et al., 2017; Li et al., 2018). This is consistent with the fact that the hand-dissected sections are composed of a large mixture of different tissues (e.g., rachis, spikelet, and floret primordia) and cell types, which in the equivalent maize ears have distinct expression profiles (Xu et al., 2021). Compared to previous bulk RNA-seq studies in developing wheat spikes, the variation observed here (measured as CV) was high among biological replicates (Supplemental Figure S2). This variation is likely caused by both biological variation (e.g., inherent variation of individual plants) and technical variation (e.g. inaccuracies in sectioning and in the developmental staging of the plant/spike) as well as the number of replicates in our analysis. A minimum of six replicates has been proposed for bulked RNA-seq (Schurch et al., 2016). Our results suggest that the RNA-seq from these small sections would benefit from a higher number of biological replicates, which should be feasible considering the high-throughput method employed for RNA extraction and library preparation, the low tissue input requirement, and the possibility to pool multiple biological replicates per sequencing lane. Despite some limitations, we could identify over 5,000 DEGs between the spatial sections for subsequent functional analysis.

In addition to G&T-Seq, several other technologies have been proposed for obtaining high resolution transcriptional profiles of plant tissues, for example, single cell RNA-seq (McFaline-Figueroa et al., 2020; Rich-Griffin et al., 2020), FACS, and the isolation of nuclei tagged in specific cell types (INTACT). These methods, however, are not spatially resolved as the complete tissue is dissolved into single cells for barcoding or selection (Rich-Griffin et al., 2020). Thus, these current methodologies do not allow, for example, to investigate whether the cell type composition of spikelets differs across the inflorescence. This would only be possible if spikelets were ‘harvested’ individually, for example through laser capture microdissection (LCM) before dissolving the tissue further into individual cells. Thiel et al. (2021) recently combined LCM followed by RNA-seq of the distinct lower/leaf ridge and upper/spikelet ridge of barley spikes. This allowed them to identify precise spatio-temporal expression patterns of many genes related to architecture and yield in barley spikes with unprecedented resolution. Looking ahead, increased resolution of Spatial Transcriptomics (currently 100 μm; (Giacomello et al., 2017)), which quantifies full transcriptomes while maintaining tissue integrity, offers the true prospect of direct localisation and quantification of gene expression. Our results argue strongly for the need of these transcriptome-wide and spatially resolved approaches to advance our biological understanding of fundamental developmental processes in plants.

### The composite nature of spikes

Early morphological studies of wheat spike development described that the stronger elongation of central spikelets during their initial establishment (glume primordia stage) first causes the lanceolate shape of the wheat spike (Bonnett, 1966). The continuous formation of primordia at the tip of the spike means that at any given growth stage, spikelets in different developmental stages will be present across the spike (Bonnett, 1966). In this study, we detected more differentially expressed genes between the three spatial sections of the spike (apical, central and basal) than between the two investigated developmental stages (Double Ridge and Glume Primordia). We identified 215 DEGs between the two developmental stages, consistent with Li et al. (2018) who identified 206 DEGs between consecutive stages across a time course of six inflorescence development stages. Feng et al. (2017) identified 753 DEGs between the Double Ridge and Floret Primordia stage, which are further apart in development than the stages used in this study. They also detected fewer DEGs when comparing early stages than between more developed spikes. By contrast, we identified 1,315 and 2,438 unique genes to be differentially expressed between the apical and basal section at DR and GP, respectively. The higher number of DEGs between spatial sections could be due to the developmental gradients occurring in the three spatial sections, which are revealed by the spatial sampling. These differences would be blurred when comparing whole inflorescences between stages due to the mixture of tissue types and spikelets at different developmental stages. A possible improvement for future transcriptome studies could be the collection of only central sections of the developing spikes or complete spatial sampling as conducted here.

The composite nature of the inflorescence tissues has been acknowledged by studies in maize (ears and tassels), where new meristems are initiated in a stepwise manner. Leiboff and Hake (2019) quantified the meristematic tissue composition of maize and sorghum tassels. For example, maize tassels in the second stage are mainly composed of spikelet pair meristems, but also contain some meristems in spikelet and inflorescence state. They concluded that the changes in these tissue compositions over time correlated well with the independently staged transcriptional changes of the tassels. Eveland et al. (2014) showed that the range of developmental ages across the maize ear, if acknowledged, can be used as an advantage in RNA-seq studies. They sequenced the tip, middle, and basal sections of 10-mm long ears independently, aiming to analyse the expression patterns in specific developmental meristem types enriched in these sections (inflorescence, spikelet, and floral meristems, respectively). The dissection of the ear therefore allowed them to study gene expression specifically for each meristematic tissue type rather than for all meristem types in intact ears. In this study, we observed that apically expressed genes are enriched for GO-terms related to “shoot system development” and “maintenance of floral organ identity”. This is consistent with the hypothesis that the apical part of the inflorescence is younger and undergoing early phases of spikelet development initiation compared to the central inflorescence section.

### Delayed transition of basal spikelets from vegetative to floral developmental programmes

We detected transcriptional gradients across the spike, with the basal section deviating most strongly from the rest of the spike. We noticed that both *SVP* and *CENTRORADIALIS (CEN)* genes remained highly expressed in the basal section of the spike, whereas their expression was lower in the central and apical sections. *In-situ* hybridisation of these genes also showed that their expression is strongest in vegetative tissue and basal spikelets in early spike development (Li et al., 2021). In contrast, *SEP1-4* and *SEP1-5* genes were expressed in the opposite gradient and showed the strongest expression in apical and central sections of the spike at Glume Primordia stage. Recent studies allow us to interpret these gradients in the context of the early steps of vegetative to floral growth transition. In wheat (Li et al., 2021; Adamski et al., 2021; Liu et al., 2021), rice (Sentoku et al., 2005; Lee et al., 2008), and barley (Trevaskis et al., 2007), *SVPs* have been characterised to be associated with vegetative growth and are downregulated upon floral transition.

In wheat, the double *SVP* mutant *vrt2svp1* leads to the formation of axillary inflorescences (Li et al., 2021). Similarly, overexpression of *CEN-D2 (TaTFL1-2D)* in wheat extends the duration of the Double Ridge stage (Wang et al., 2017), whereas loss-of-function mutations in barley *CEN* suggest they repress floral development under short-day conditions (Bi et al., 2019). Double knockout mutants of the MADS-box *SQUAMOSA* genes *vrn1ful2* highlighted that these two genes act as transcriptional repressors of *SVP* and *CEN* genes in early wheat spike development (Li et al., 2019). Furthermore, through a series of genetic and biochemical studies, Li et al. (2021) showed that the downregulation of *SVP* genes is necessary for the formation of flowering promoting MADS-box protein complexes including VRN1, FUL2 and SEP proteins. Hence the coordinated downregulation of *SVPs*, and possibly *CEN* genes, along with the upregulation of *SEP* genes is required for normal floral transition and spikelet development in wheat. Previous studies in rice have found similar expression patterns, as well as mutant effects, of *SVPs* and *SEPs* suggesting a conserved function in flowering transition across the two species (Ren et al., 2016; Wu et al., 2018).

Based on our results, the floral developmental programme across the wheat spike appears to be most advanced in its apical and central sections, while being delayed in the basal sections. We hypothesise that this is due to elevated *VRT2* expression at the base of the spike, which hinders the progression of the flowering programme via *SEP* class flowering genes. Likewise, the higher expression levels of the wheat *CEN2* and *CEN5* homologs at the base are consistent with a delay in floral transition that could interfere with the development of the spikelet primordia. Therefore, although the basal spikelet primordia are initiated first chronologically, their developmental age in terms of the floral programme is delayed with respect to the more recently formed central and apical spikelet primordia. This could explain in part why the spikelet primordia in the basal region of the spike elongate less and develop slower than central spikelets despite being initiated first (Bonnett, 1966). Likewise, the less advanced floral developmental programme could also explain why the overexpression of *SVPs* in barley (*HvBM10*) leads to complete floral reversion in basal but not apical spikelets (Trevaskis et al., 2007).

We hypothesise that *SVPs* need to be downregulated upon floral transition to allow timely establishment and progression of the early spikelet primordia. Failure to do so would delay their development and result in their final rudimentary shape in the mature spike. In line with this hypothesis, we observed increased RBS in genotypes with prolonged and increased *VRT2* expression in a dosage-dependent manner. In our qRT-PCR data, we also observe increased expression of *SVPs* alongside reduced expression of *SEPs* in *VRT-A2b* lines at Double Ridge and Glume Primordia stage. However, we cannot exclude the possibility that the increase in *VRT2* expression in the *VRT-A2b* lines could also affect basal spikelet development at a later stage of spike developmental (e.g., from Terminal Spikelet stage to anthesis). We are currently using quantitative live imaging to compare cellular growth dynamics of spikelets at different stages of spike development between the *VRT2* NILs.

The finding that the expected downregulation of *SVPs* and *CENs* does not follow the chronological age of the tissues suggests that other gradients across the spike might influence spikelet development. Debernardi et al. (2017) showed that in tetraploid wheat *AP2-5* and miR172 have consistent and opposing expression gradients across the spike at three consecutive developmental stages. The persistent expression gradient of *AP2-5* supports the idea that expression patterns across the spike, beyond the ones caused by age differences of spikelets, exist. Furthermore, they proposed a model illustrating that the phenotypic effect of mutants across the spike differs due to the existing gradient of expression of this gene (Debernardi et al., 2017). Other examples of mutants with different phenotypic effects across an inflorescence include *vrn1ful2* (Li et al., 2019) in wheat, *tassel sheath1* (*tsh1*, (Whipple et al., 2010)) and *ramosa2 (ra2*, (Bortiri et al., 2006)) in maize, *SEPALLATA* double mutant *Osmads5Osmads34* in rice (Zhu et al. 2021), as well as *many noded dwarf1* (*mnd1*, (Walla et al., 2020)), *frizzy panicle* (Poursarebani et al., 2015), and *vrs2* in barley, which was also found to be consistently differentially expressed across the spike (Youssef et al., 2016). *VRS2* has been shown to maintain a basal to apical expression pattern across three, post awn initiation developmental timepoints in barley (Youssef et al., 2016). The study of *vrs2* mutants revealed that *VRS2* is furthermore engaged with the basal-apical patterns of auxin, cytokinin, and gibberellin across the spike. While hormonal gradients across the spike in early development have not been studied in great detail in wheat, they have been shown to play crucial roles in floral induction and development in *Arabidopsis* (Reinhardt et al., 2000). Their patterns across the spike should be investigated in future studies addressing developmental differences across the spike.

### A model for the regulation of leaf and spikelet ridge outgrowth in the base of the spike

Recently, Meir et al. (2021) proposed that in shoot apical meristems of tomato, similar to processes during embryonic development, transient programmes are required to inhibit a preceding setup (i.e. vegetative growth), before a new developmental program (flowering) can be initiated. We propose that the altered gene expression and development of the basal spikelets could be a consequence of their initiation during the transient phase between vegetative and floral network shifts and thus being exposed to mixed signals of development. Upon floral transition, the lower (leaf) ridge is supressed, while the growth of spikelet ridges from the previously supressed axillary meristems is activated. Development of lower ridges subtending all branching events is suppressed in grass inflorescences upon flowering transition (Whipple et al., 2010). Li et al. (2019) noticed that this suppression was disrupted in the double *vrn1ful2* and triple *vrn1ful2ful3*-null mutants, which fail to down-regulate *SVP* genes. In these mutants, the upper spikelet meristems generate vegetative structures resembling tillers that are subtended by bracts or leaves originating from the lower leaf ridge.

We observed that genes that were highly expressed in the basal section of the inflorescence (cluster 4) have previously been shown to be expressed specifically in the lower/bract ridge and before or at vegetative to floral transition. This is also supported by the GO-term enrichment of photosynthesis related terms in cluster 4. Our tissue sections do not allow us to distinguish lower and upper ridge tissues, however, the two ridges have been separately collected and sequenced via LCM in barley (Thiel et al., 2021). In this barley dataset, we found a higher expression of *HvVRT2* (*HORVU7Hr1G036130*) and *FLOWERING LOCUS C* (*HvFLC; HORVU7Hr1G054320*) in the lower ridge compared to the upper ridge, whereas *HvSVP1* (*HORVU6Hr1G077300*) was also marginally more highly expressed (Supplemental Figure S5). Furthermore, the barley *MND1* gene (*HORVU7Hr1G113480*) has recently been shown to be expressed in leaf primordia and during the Double Ridge stage in the basal region of the spike in barley (Walla et al., 2021), while it is most highly expressed in the vegetative meristem and lower/leaf ridge in the LCM data (Thiel et al 2021). We observed that in our data, the wheat *MND1* orthologs (*TraesCS7A02G506400, TraesCS7B02G413900, TraesCS7D02G494500*) were significantly more highly expressed in the basal section than the apical section at both DR and GP stage (Supplemental Table S2). The suppressed leaf ridge (or bract) has been proposed to act as a signalling centre, regulating the fate of the upper spikelet meristem ridge (Whipple, 2017). Insufficient bract suppression during the formation of the basal spikelets might therefore negatively affect initiation and development of spikelets.

At DR, the widest point of the spike is indeed as expected the base and not the central section (Figure 1A). The lower ridge is however much less developed in the central section and can be hardly seen in the apical ridges. Interestingly, mutants failing to repress the lower ridge growth, such as *third outer glume1* (*trd1*), the barley ortholog of maize *tsh1*, develop large bracts from the lower ridge in basal spikelets, unlike apical spikelets, which do not develop bracts from their lower ridges regardless of the absence of *TRD*. This is reminiscent of the gradient in the strength of the phenotypic effects observed from the top to the base of the inflorescence in multiple *Poaceae* mutants (discussed above). We therefore hypothesise that the basal meristems develop into smaller spikelets and larger bract primordia due to a slow suppression of “vegetative growth signals” (e.g., *SVPs*) and a concomitant slow upregulation of “floral growth signals” (e.g., *SEPs*) upon floral transition. To investigate how a change from vegetative to floral signalling might affect the development of individual meristems, we modelled the genetic interaction of *SVPs* and *SEPs*, as proposed by Li et al. (2021), in the spatial context of a growing spike (Supplemental File S1). Under the assumptions that SVP supresses *SEP* expression, *SVP* expression is downregulated upon flowering, and that SEP promotes spikelet outgrowth, the model could recapitulate (a) the observed opposing gradients in expression of *SVPs* and *SEPs* along the spike, and (b) the formation of a lanceolate shaped wheat spike with reduced spikelet elongation and stronger bract growth in the most basal spikelets. Thus, whilst this hypothesis will require further investigation and testing, modelling supports its plausibility.

## Materials and Methods

### Plant materials

Hexaploid wheat (*Triticum aestivum*) germplasm used in this study includes wildtype hexaploid wheat cultivar Paragon and *P1/VRT2 germplasm* described in Adamski et al. (2021) including *P1* NILs, recombinants, and T_1_ transgenic lines carrying the *T. polonicum VRT-A2b* copy under the native promoter. *T. polonicum* accession T1100002 was obtained from the John Innes Centre Germplasm Resources Unit (https://www.seedstor.ac.uk/search-infoaccession.php?idPlant=27422). For field experiments, we used between two to four sibling BC_4_/ BC_6_ NILs differing for the *VRT-A2b* allele.

### Low input RNA sequencing

Paragon seedlings were grown in a single batch in a controlled environment growth chambers in 24-cell seed trays under long-day (16 h light/8 h dark) photoperiods at 300 μmol m^-2^ s^-1^, with a day temperature of 20 °C and a night temperature of 15 °C. Inflorescences for Double Ridge (DR) stage were collected 18 days after sowing, while inflorescences for Glume Primordia (GP) stage were collected 22 days after sowing. All plants were grown in “John Innes Cereal Mix” (40% Medium Grade Peat, 40% Sterilized Soil, 20% Horticultural Grit, 1.3 kg·m^-3^ PG Mix 14-16-18 +Te Base Fertiliser, 1 kg·m^-3^ Osmocote Mini 16-8-11 2 mg + Te 0.02% B, Wetting Agent, 3 kg·m^-3^ Maglime, 300 g·m^-3^ Exemptor).

Four individual spikes per developmental stage (DR and GP) were dissected into apical, central, and basal sections (1:1:1 ratio) using a stereo microscope (Leica MZ16). Sections were immediately placed into 96-well plates (on ice) containing 10 μL of RLT plus (Qiagen, Hilden, Germany). All instruments and surfaces were cleaned with 80% ethanol, RNAse-free water and lastly RNAse-out solution after each sample to reduce cross-contamination and RNA degradation. Samples were stored at −80 °C until cDNA preparation, using the G&T-seq method as previously described (Macaulay et al., 2015). cDNA was normalised to 0.2 ng/ μL before Nextera (Illumina, San Diego, CA, USA) library preparation using a Mosquito HV liquid handler (STP, Royston, UK) in a total reaction volume of 4 μL as described in Mora-Castilla et al. (2016). Libraries were pooled by volume and sequenced on a single lane of a NovaSeq 6000 (NVS200S2 flow cell, 100 bp paired-end reads).

### Bioinformatic analysis

For the RNA-seq analysis, we used the RefSeqv1.0 genome assembly and the RefSeqv1.1 gene annotation (https://urgi.versailles.inra.fr/download/iwgsc/IWGSC_RefSeq_Assemblies/v1.0/; IWGSC et al. (2018)). Reads were trimmed and adapters were removed using trim-galore v.0.4.2 (https://www.bioinformatics.babraham.ac.uk/projects/trim_galore/) with settings: “--paired --fastqc --a GGTATCAACGCAGAGT --clip_R1 20 --clip_R2 20 --trim-n”. Minimum length of reads retained was set to 50 bp. Reads were aligned to the RefSeqv1.0 genome assembly using HISAT2 v. 2.1.0 (https://daehwankimlab.github.io/hisat2/; Kim et al. (2019)) with the following parameters: “--pen-noncansplice 20 --mp 1,0 --rna-strandness RF”. Alignment files were converted to BAM format, sorted, indexed, filtered, and purged of all none-primary alignments (0×100 flag) using samtools (v. 1.9; Li et al. (2009)). HTSeq v.0.6.1 (https://htseq.readthedocs.io/en/master/; Anders et al. (2015)) was used to count the read numbers mapped to the RefSeqv1.1 gene models.

HT-read count normalization and differential expression analyses were performed using the DESeq2 v. 1.28.1 R packages (https://bioconductor.org/packages/release/bioc/html/DESeq2.html; Love et al., 2014; RStudio 1.2.5001). Genes with an average expression below 10 HT-count, and which were not expressed (i.e. ≤ 10 HT counts) in at least three libraries, were removed from the analysis. Correlation between expressed genes and Waddington stage and/or section was tested by ANOVA. Raw read data from Li et al. (2018) and Feng et al. (2017) were pseudo-aligned using Kallisto Sleuth pipeline (https://scilifelab.github.io/courses/rnaseq/labs/kallisto) and the coefficient of variation was calculated for each gene (by condition) using R (RStudio 1.2.5001) ddply (plyr 1.8.6). Differentially expressed genes (DEGs) between the two Waddington stages (DR and GP) were calculated with the design “~plant + section”, while DEGs between the three sections (apical, central, basal) were determined using the design “~section + waddington:plant + waddington”. DEGs among the sections within each Waddington stage were determined with the design “~plant + section”. For each gene, an adjusted *P*-value was computed by DESeq2 (using the using the Benjamini and Hochberg method (Benjamini and Hochberg, 1995)), and those with an adjusted *P*-value of ≤ 0.05 were considered differentially expressed. DeSeq2 also computed Log2FoldChanges as well as the associated uncertainty (lfcSE, see Love et al. (2014) for further detail). The “contrast” function was used to determine pairwise comparison *P*-values. The full set of expression data and comparisons is presented in Supplemental Data Set S1. Enrichment of GO-terms was performed using the online tool “PLAZA” (https://bioinformatics.psb.ugent.be/plaza; Van Bel et al. (2017)) using the recommended settings, and all enriched GO-terms of Biological function (BF) and Cellular Compartment (CC) were retained. In brief, PLAZA determines the overrepresentation of a certain GO-term in a gene set compared to the genome-wide background frequency (= all expressed genes in this experiment; submitted manually). The significance of over- or underrepresentation is determined using the hypergeometric distribution and the Bonferroni method is applied to correct for multiple testing. Note that enrichment folds are reported in log2 fold scale. Enrichment of TF families (Genes that were annotated as TFs were obtained from https://opendata.earlham.ac.uk/wheat/under_license/toronto/Ramirez-Gonzalez_etal_2018-06025-Transcriptome-Landscape/data/data_tables/ (Ramirez-Gonzalez et al., 2018)) and MADS-box TFs (based on Schilling et al. (2020)) was performed in R using the phyper() function from stats package v.4.0.1 to test for Hypergeometric Distribution. All DEGs were scaled and centred using R-base function “scale”. All cluster analysis was performed on scaled data using R (stats) functions kmeans and hclust, followed by visualisation through pheatmap v. 1.0.12 (https://cran.rstudio.com/web/packages/pheatmap/index.html). Correlation to centroid cluster shape of each gene expression pattern was calculated using the “cor” function from R stats.

### Quantitative real-time PCR analysis

*P1* NILs were grown in controlled growth chambers in 24-cell seed trays under the same conditions as used in the low input RNA-seq experiment (see above). For each biological replicate, we pooled 30 inflorescences for DR stage, 15 for GP stage, and nine for Terminal Spikelet stage (n = 4 biological replicates per stage). Inflorescences from NILs were dissected using a stereo microscope (Leica MZ16). Inflorescences were dissected into apical, central, basal and peduncle sections (1:1:1:1 ratio). At Double Ridge stage, inflorescences were only dissected into apical, basal and peduncle section as the inflorescences were too small to be accurately dissected into four sections for all 30 plants per biological replicate. Each section was immediately placed into 1.5-mL tubes on dry ice and tubes were snap frozen in liquid nitrogen as soon as all plants for the sample were collected. Samples were stored at −80°C until needed. Inflorescences were collected within 2-3 hours, 9 hours after the lights came on in the growth chamber. Tissue was homogenized in a TissueLyser II (Cat No.: 85300, QIAGEN) using 3-mm steel beads (Cat No.: 69997, Qiagen); tubes were shaken for 20-s at 28 Hz with dry ice.

All RNA extractions were performed using the RNeasy Plant Mini Kit (Cat No.: 74904, Qiagen) with RLT buffer according to the manufacturer’s protocol followed by RNA ethanol precipitation (https://projects.iq.harvard.edu/files/hlalab/files/ethanol-precipitation-of-rna_hla.pdf). DNA digestion was performed using the RQ1 RNase-free DNase set (Cat No.: M6101, Promega) according to the manufacturer’s protocol. RNA was reverse transcribed using M-MLV reverse transcriptase (Cat No.: 28025013, Thermofisher) according to the manufacturer’s protocol. For the qRT-PCR reactions, LightCycler 480 SYBR Green I Master Mix (Roche Applied Science, UK) was used according to the manufacturer’s protocol. The reactions were run in a LightCycler 480 instrument (Roche Applied Science, UK) under the following conditions: 10 min at 95 °C; 40 cycles of 10 sec at 95 °C, 15 sec at 62 °C, 30 sec at 72 °C; dissociation curve from 60 °C to 95 °C to confirm primer specificity. All reactions were performed with three technical replicates per sample and using *TaActin* as the reference gene (Uauy et al., 2006). Relative gene expression was calculated using the 2^-ΔΔCt^ method (Livak and Schmittgen, 2001) with a common calibrator so that values are comparable across genes, tissues, and developmental stages. All primers used in qRT-PCR came from Adamski et al., 2021 can be found in Supplemental Table S10.

All qRT-PCR data was normalised using a log2 transformation. A three-way ANOVA including Waddington stage, section, and genotype yielded significant two-way interactions. The differences between sections of the genotypes were therefore further analysed individually for each Waddington stage and genotype. For each of the two genotypes we individually performed Tukey multiple comparison tests to determine differences between the sections within each developmental stage by Tukey multiple comparison test. Differences between the two genotypes were also analysed individually for each Waddington stage. Furthermore, the differences between the genotypes were investigated individually for each section within the Waddington stage if the interaction term was significant (in GP and TS). For all analysis see Supplemental Table S6.

### Field experiments and phenotyping

*VRT-A2* NILs were evaluated in four field experiments. Three trials were located at The Morley Agricultural Foundation trials site, Morley St Botolph, UK (52°33’15.1”N 1°01’59.2”E) in 2017, 2018 and 2020 and one trial was sown in 2020 at the John Innes Experimental trials site in Norwich, UK (52°37’50.7”N 1°10’39.7”E). In Morley (2017) we analysed two BC_4_ lines of *VRT-A2a* and three BC_4_ lines of *VRT-A2b*. In Morley (2019) we analysed two BC_6_ and one BC_4_ line per *VRT-A2* allele and in Morley and Church Farm 2020 we analysed two BC_6_ and two BC_4_ lines for each *VRT-A2* allele. All experiments were drilled as yield-scale plots (6 m x 1.2 m) and sown by grain number for comparable plant densities aiming for 275 seeds m^-2^. The trials were arranged in a randomised complete block design (RCBD) with five replicates per sibling line per location. Developmental and plant architecture traits were evaluated throughout the growing period. A 10-ear grab sample was collected from each plot pre-harvest for the assessment of rudimentary basal spikelet (RBS) numbers and other phenotypes (recorded in Adamski et al. 2021). RBS were defined as spikelets carrying no grain at maturity and counted for each spike individually. To determine the differences between the *P1^POL^* and *P1^WT^* NILs, we performed analysis of variance (ANOVA) on the multiple field trials phenotypic data. For the analysis of individual trials, we used a two-way ANOVA including Genotype + Block performed in R (‘car’ package version 3.0-10; RStudio 1.2.5001).

### Glasshouse phenotyping

We evaluated the BC_4_ NILs and BC_6_F_3_ recombinant lines, as well as *T. polonicum* accession T1100002, under standard glasshouse conditions. 18-20 plants per genotype were grown in 1 L pots containing John Innes Cereal Mix under long day conditions (16 h light, 8 h dark). The genotypes of all plants were confirmed using KASP marker *SP1Pol* (Adamski et al., 2021).We counted the number of rudimentary basal spikelets (RBS) for all tillers of all biological replicates at maturity. To evaluate the differences in RBS between genotypes, we performed a two-way ANOVA analysis and post-hoc multi-pairwise comparisons Sidak test (‘car’ package version 3.0-10; RStudio 1.2.5001).

### Phenotyping of transgenic lines

T_1_ lines from Adamski et al. (2021) differing for the copy number of the *VRT-A2b* transgenic construct (zero = 0 copies; low = 1-5 copies; high = 9-35 copies) were grown in 1 L pots with John Innes Cereal Mix under 16 h light at 20°C and 8 h dark at 15°C in controlled environment growth chambers. We measured RBS number for the main tiller of all plants at maturity. To determine differences in RBS between the three transgenic classes, we performed analysis of variance (two-way ANOVA; ‘car’ package version 3.0-10). We performed Dunnett tests to compare the low and high copy lines against the zero copy number controls (RStudio 1.2.5001).

### Modelling

The computational model of wheat spike shape formation was developed using the multi-agent programming language and modelling environment, Netlogo (Wilensky, 1999). Gene interactions were modelled as previously described (Li et al., 2021). The model can be accessed via the interactive web-version of the model (Supplemental File S1).

In brief, both spikelets and leaves are initiated with rates that depend on the levels of SEP. Leaf initiation rates are suppressed by SEP, whereas spikelet initiation requires SEP. The maximum initiation rates are the same for both spikelets and leaves but different before (*r_vegetative_*) and after (*r_flowering_*) flowering. Once initiated, the leaves and spikelets grow at a rate defined by the parameters *r_leaf_* and *r_spikelet_*, respectively. Leaf growth does not depend on SVP or SEP levels, whereas spikelets only increase in size every iteration if their SEP level is above a given threshold (*SEP_growth_threshold_*). Expression of both *SVP* and *SEP* only occurs at meristem initiation. After this, the levels of SVP and SEP cannot increase, although SEP is degraded. SVP is not degraded, solely because at this point, nothing is dependent on SVP levels, whilst spikelet growth depends upon SEP levels.

*SVP* expression rates start to decrease, once flowering is triggered, according to:

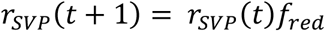

where *r_SVP_*(*t*) is the rate of *SVP* expression at that time step, and *f_red_* is a rate reduction factor. *SEP* expression depends upon the levels of SVP in the meristem in which the initiation points are located, depending on a Hill function (Alon, 2007),

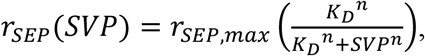

where *r_SEP,max_* is the maximum rate of *SEP* expression, *K_D_* is the binding constant, and *n* is the Hill coefficient. The resulting curves for *SVP* and *SEP* expression are shown in Supplemental Figure S6.

SVP levels are initiated with the current value of *r_SVP_*. SEP levels are initiated using *r_SEP_*, and reduce by degradation rate, *δ_SEP_*, following

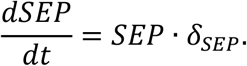

## Supporting information

Supplementary Data Set

NetLogo Model

## Data Availability

The raw RNA-seq read libraries used in this study are available from NCBI BioProject PRJNA749586.

## Acknowledgments

We thank Phil Robinson from the Scientific Photography at JIC for taking high-quality images of spikes and spikelets and Tobin Florio (http://flozbox-science.com/) for scientific illustrations for Figure 1. We thank the JIC Field Experimentation and Horticultural Services teams for technical support in field and glasshouse experiments and the JIC Bioimaging facility and staff for their services and technical support.

## Additional Files

**Supplemental Figure S1:** Principal Component analysis (PCA) of RNA-seq libraries.

**Supplemental Figure S2:** Comparison of Coefficient of variation (CV) of gene expression in wheat RNA-seq data sets.

**Supplemental Figure S3:** Expression patterns of al DEGs clusters (n=7) as identified by k-means clustering.

**Supplemental Figure S4:** Dissected Floret 1 and 2 of basal and central spikelets of BC_6_ NILs before anthesis.

**Supplemental Figure S5:** Expression of barley genes in Thiel et al. (2021), which are orthologous to wheat genes highly expressed in basal spike sections.

**Supplemental Figure S6:** Example of Netlogo simulation outcome with default parameters.

**Supplemental Table S1:** Quality control measurements of all 24 RNA-seq libraries.

**Supplemental Table S2:** Summary of normalised gene expression, statistical analyses and k-means clustering for all differentially expressed genes (*Padj* ≤ 0.05).

**Supplemental Table S3:** Enrichment of Gene Ontology (GO)-terms in the seven identified clusters of DEGs.

**Supplemental Table S4:** Enrichment of Transcription factor families and MADS-box transcription factor genes in the seven identified clusters of DEGs.

**Supplemental Table S5:** Relative expression of *VRT-A2* and *SEP1-4* measured in Paragon NILs with either the wildtype (*VRT-A2a*) or *T. polonicum* allele (*VRT-A2b*).

**Supplemental Table S6:** Statistical analysis of qRT-PCR data from *VRT-A2* NIL spike sections (Supplemental Table S5).

**Supplemental Table S7**: Field evaluations for rudimentary basal spikelets (RBS) in *VRT-A2* NILs.

**Supplemental Table S8**: Graphical genotype from Adamski et al (2021) and RBS phenotype of BC_6_ recombinant inbred lines (RILs) for *VRT-A2*.

**Supplemental Table S9**: Rudimentary basal spikelet phenotypic data from *VRT-A2* transgenic lines.

**Supplemental Table S10:** List of primers used in qRT-PCR in this study.

**Supplemental Dataset S1:** Summary of normalised gene expression, statistical analyses and k-means clustering for all expressed genes.

**Supplemental File S1:** Interactive web-version of the wheat spike model.

