## Supplementary Figures for "High expression of *VRT2* increases the number of rudimentary basal spikelets in wheat"

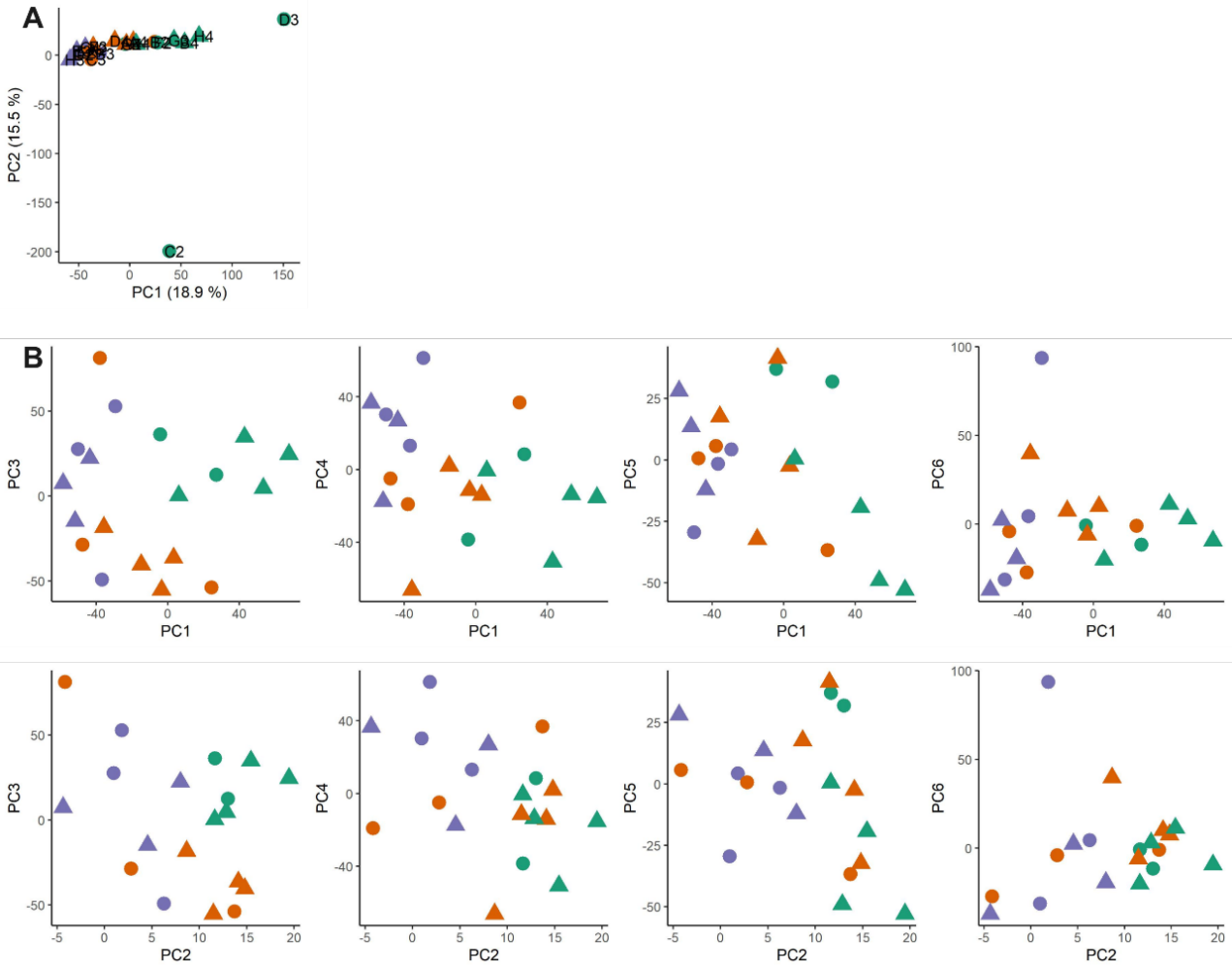

**Supplemental Figure S1:** (A) Principal Component (PC) analysis of 21 libraries that passed initial quality control (three libraries excluded due to low average number of expressed genes), indicating two outlier libraries (C2&D3) which were subsequently removed from the analysis. (B) PCs of the 19 libraries that passed all controls. Purple = Apical, Orange = Central, Green = Basal; Circle = Double Ridge, Triangle = Glume Primordia stage

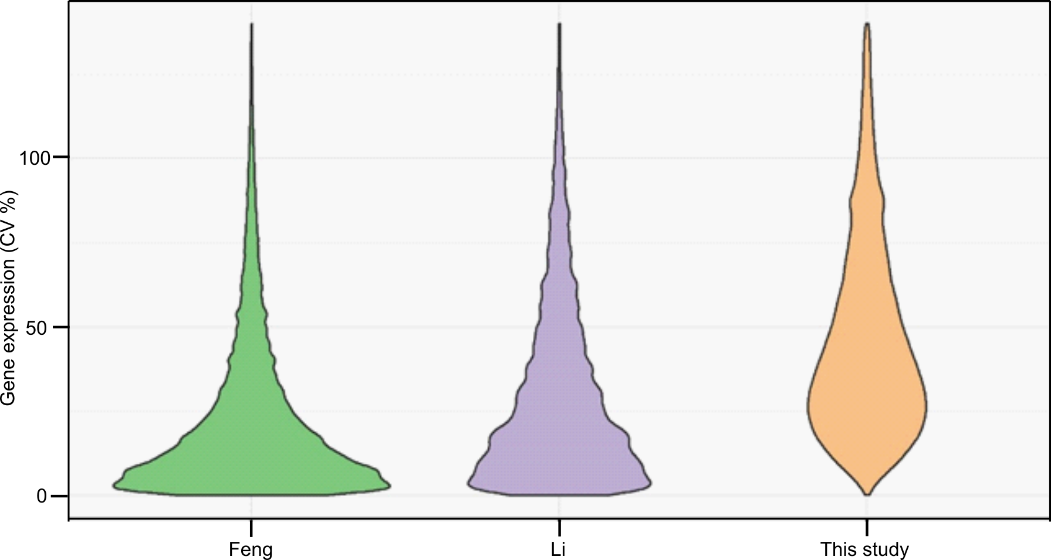

**Supplemental Figure S2:** Coefficient of variation (CV) of gene expression. CVs among the biological replicates were calculated for all expressed genes in this study (number of biological replicates = 2-4). Feng *et al*. (2017) and Li *et al*. (2018) have n=2 biological replicates of pooled samples.

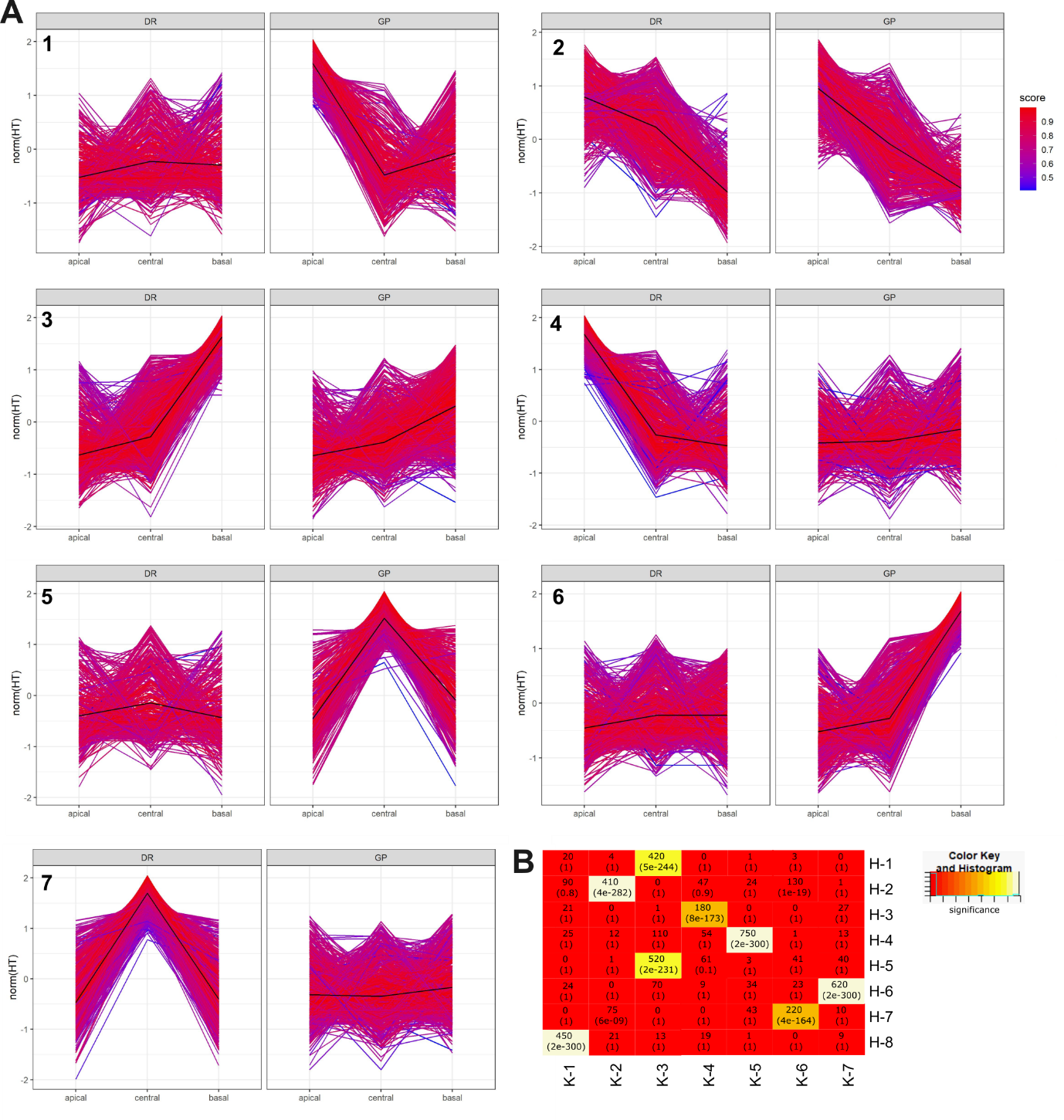

**Supplemental Figure S3:** (A) Expression pattern of differentially expressed genes (DEGs) in the seven clusters identified by k-means clustering. Score colouring indicates the correlation of gene expression with the centroid pattern of the cluster (black line). Expression values were normalised using DeSeq2, scaled and centred (R(base) function “scale”;see Methods). (B) Correlation of k-means (K) and hierarchical (H) clusters. Upper value is the number of shared genes, lower value is the P-value of the correlation. Calculation was performed using WGCNA “overlapTable” function (Langfelder & Horvath, 2008). Colours = -log(P-value) (red=less significant, white = most significant).

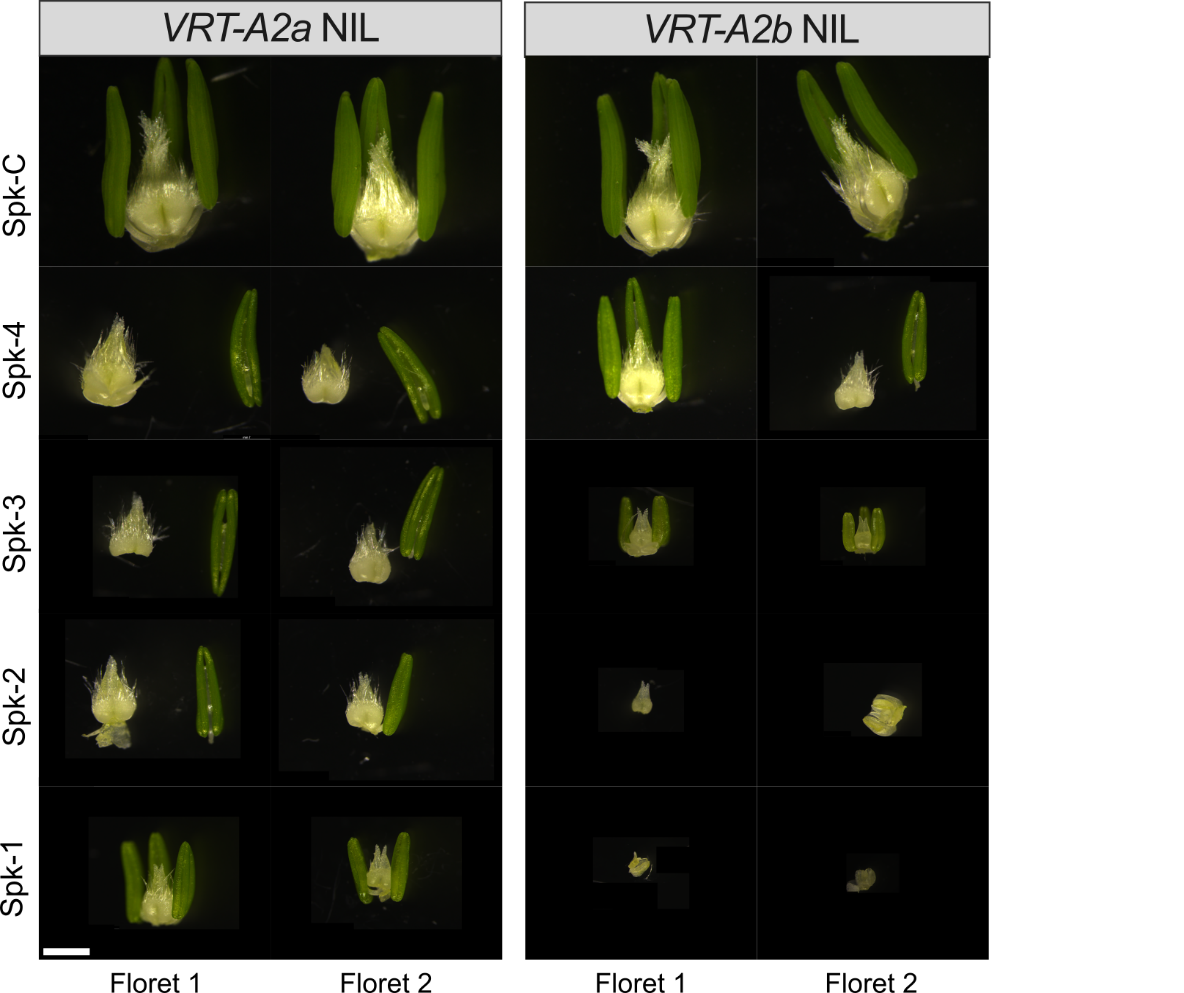

**Supplemental Figure S4:** Dissected Floret 1 and 2 of basal and central spikelets of BC_6_ NILs approximately 1 week before anthesis. We dissected florets 1 and 2 of the basal four spikelets (Spk-1 to Spk-4) as well as the floret of the biggest central spikelet (Spk-C). NILs were grown in glasshouse conditions (2021). Scale = 1mm. Note that not all pictures include all three anthers.

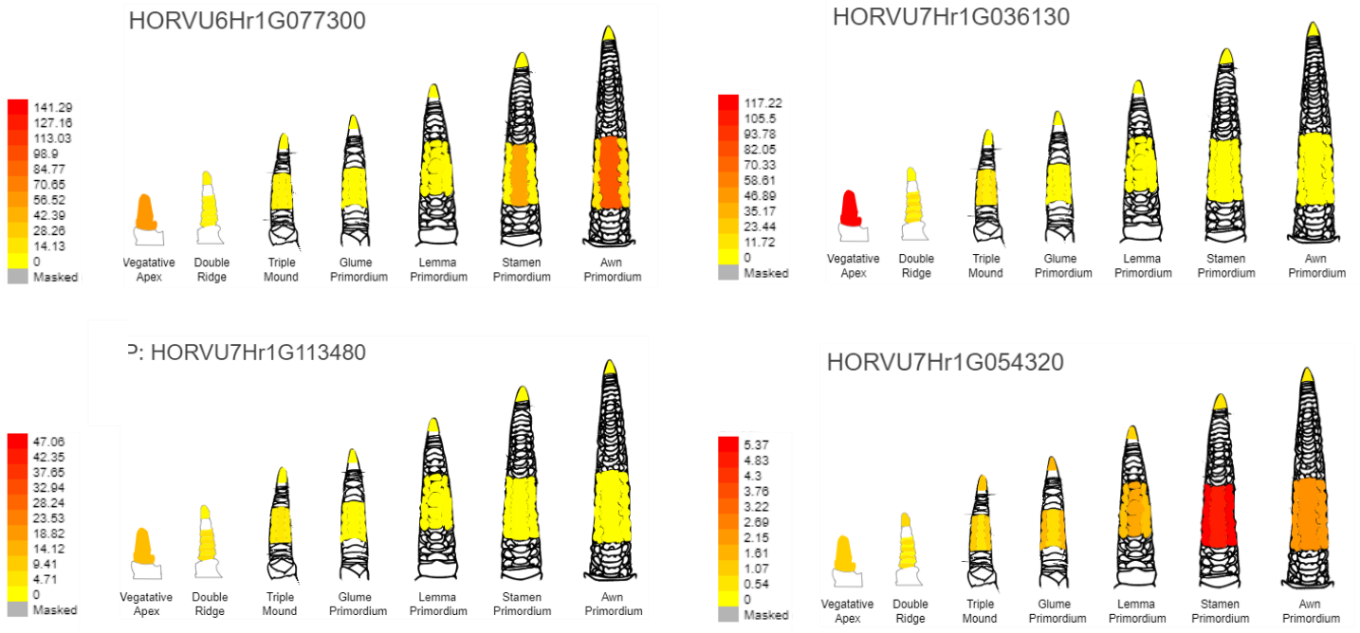

**Supplemental Figure S5:** Expression of barley genes *HvSVP1* (*HORVU6Hr1G077300*), *HvVRT2* (*HORVU7Hr1G036130*), *HvMND1* (*HORVU7Hr1G113480*), and *HvFLC* (*HORVU7Hr1G054320*), which are orthologous to wheat genes that are highly expressed in basal spike sections. The barley orthologs are expressed more strongly in the lower leaf ridge than the upper. Data from Thiel *et al.* (2021) and visualized in the barley eFP gene expression browser (<http://bar.utoronto.ca/>).

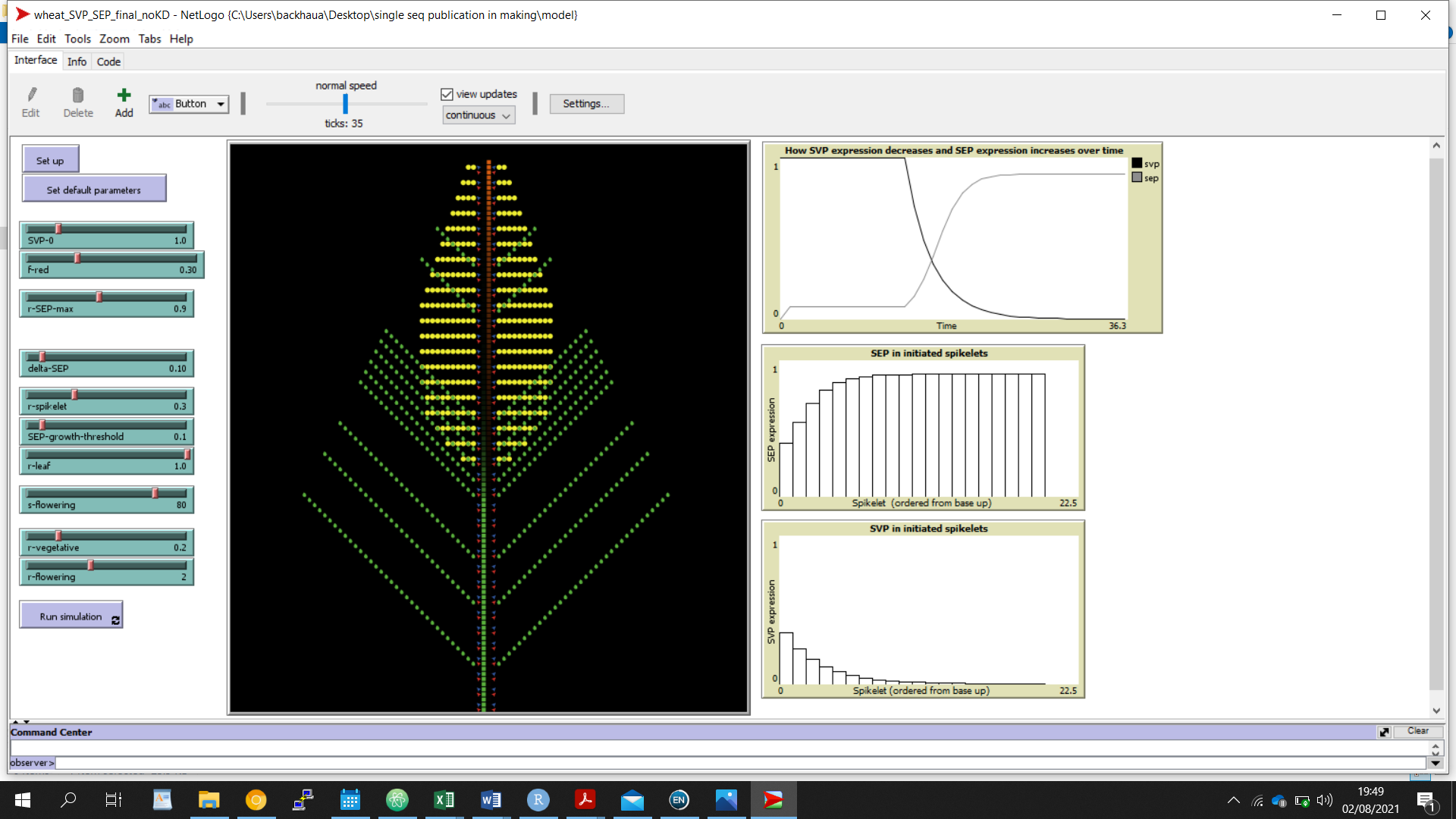

**Supplemental Figure S6:** Example image of the Netlogo model of wheat spike development. Green = leaf tissue, Yellow = spike tissue, red = meristem initiation point committed to leaf development. Blue = axillary meristem committed to spikelet/branch. The model uses expression of *SEP* and *SVP* class genes to predict when meristems (red) produce leaf tissue (green) and when they switch to producing spike tissue (yellow). See methods for model parameters and use. Top-right hand graph shows the decrease of *SVP* expression and the increase of *SEP* expression over time. The middle and bottom graphs depict the gradients of *SEP* and *SVP* expression, respectively, from the basal to the apical spikelets. The modelled growth (central graph) shows how basal spikelets are smaller than their central counterparts and have leaf ridge growth due to the higher *SVP* and lower *SEP* expression.
